# Differing drivers of range-wide genetic diversity across previously glaciated Northern hemisphere landscapes

**DOI:** 10.1101/2025.04.28.650964

**Authors:** Lu Liu, James S. Borrell, Nian Wang

## Abstract

- The latitudinal gradient (LG) hypothesis predicts a poleward decrease of genetic diversity. Similarly, the central-marginal hypothesis (CMH) predicts higher genetic diversity in range centres than margins. We examined these patterns in Europe (EU), North America (NA) and East Asia (EA), which experienced contrasting patterns of landscape fragmentation.
- We compiled genetic variation data for 445 plant species and 8,530 populations. We calculated ecological metrics comprising distance to the range margin and native climatic niche margin, distance to refugia, latitude and altitude. We applied Bayesian inference to evaluate the relationships between genetic variation and ecological metrics.
- We found universal support for the CMH, with populations at the centres of species ranges and/or native climatic niches exhibiting higher genetic diversity than those at the margins and stronger patterns in woody versus herbaceous plants. Evidence for a LG in diversity was less consistent with latitude: negatively correlated with genetic diversity in EU, positively correlated in EA, and no correlation in NA.
- Our analysis supports the view that the LG is largely driven by latitudinal trends in glaciation, particularly in EU, but highlights that glaciation patterns and their resulting impacts on landscape genetic diversity were more heterogeneous in NA and EA.

## Introduction

Contemporary patterns of genetic variation are strongly shaped by the complex interactions of historical climate and biogeography. Two hypotheses have been developed to explain widespread observational patterns in genetic diversity across landscapes. First, the latitudinal gradient (LG) hypothesis proposes that genetic diversity should mirror patterns in species diversity and be higher at lower latitudes (**Fig. 1a**) (Pereira, 2016). This is because during the last glacial maximum (LGM), species survived through latitudinal or altitudinal range contraction and expansion (Hewitt, 2000; Shafer *et al*., 2010; Qiu *et al*., 2011; Liu *et al*., 2012). These processes eroded the effective population sizes, attenuating genetic variation at higher latitude. The LG in genetic diversity has been illustrated in several global studies (Gratton *et al*., 2017; Fonseca *et al*., 2023), but more nuanced observations have also demonstrated higher diversity at intermediate latitudes due to admixture (Petit *et al*., 2003).

**Fig. 1.**
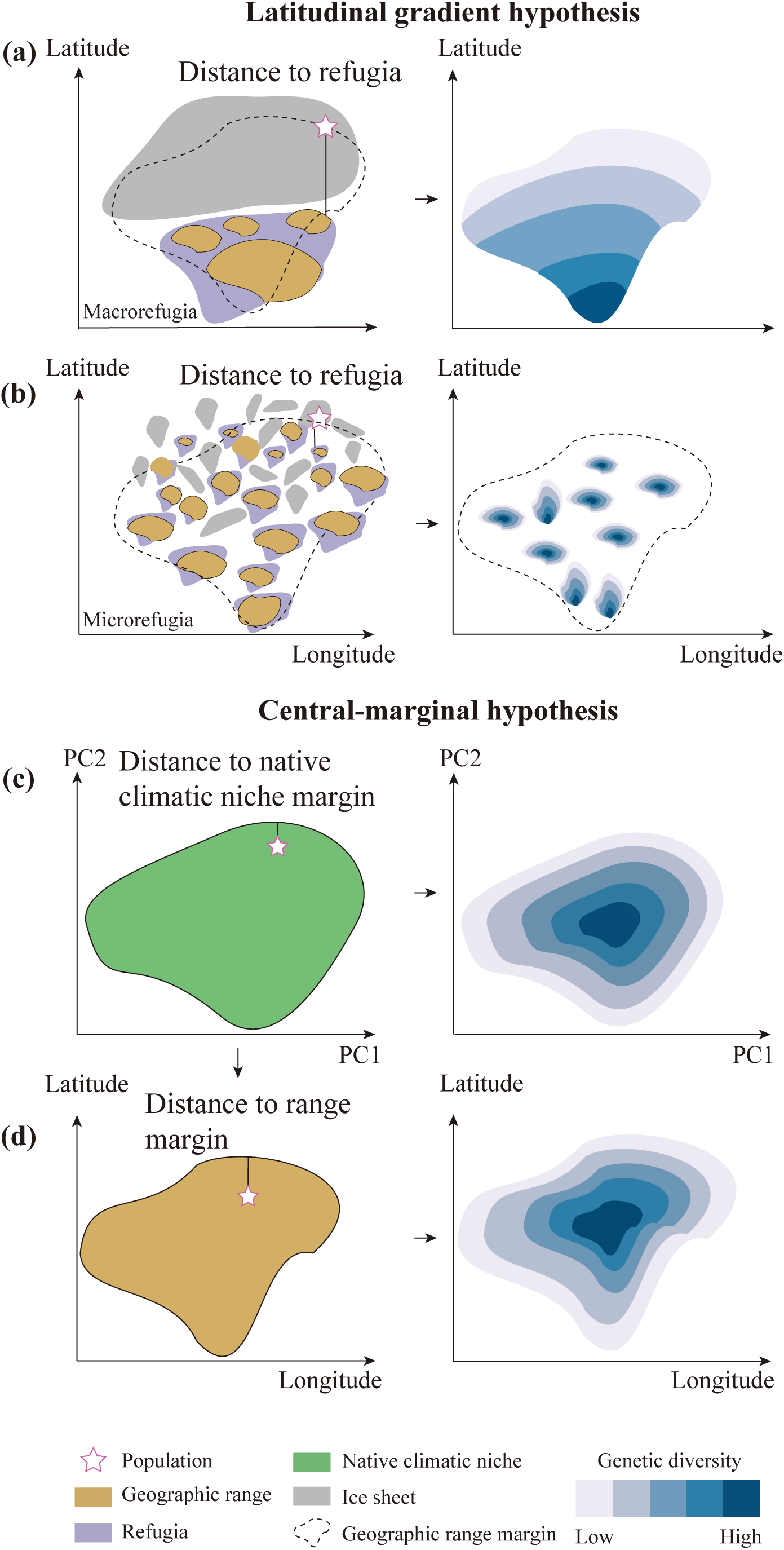
A conceptual framework illustrating the influence of climate and geography on population genetic diversity. (a, b) The latitudinal gradient and resulting patterns of genetic diversity under continuous and fragmented icesheets. (c, d) the expected patterns of genetic diversity under the central-marginal hypothesis.

Second, the central-marginal hypothesis (CMH) posits that populations at the species’ range margin exhibit lower genetic diversity compared to those at the range’s core (**Fig. 1d**) (Eckert *et al*., 2008; Pironon *et al*., 2017). The CMH predictions are based on an assumed concordance between species niche and range position such that optimal ecological conditions peak at the centre of a species’ range and decline towards the periphery (Pironon *et al*., 2017). Consequently, populations at the range margins experience greater environmental fragmentation and lower climate suitability compared to those at the range centre, hence are more vulnerable to loss of genetic variation due to genetic drift (Kawecki, 2008; Polechová & Barton, 2015). The magnitude of glaciation is likely to impact the severity of range shifts and subsequent landscape genetic diversity pattners arising from the CMH.

While expansion of Northern Hemisphere ice sheets was widespread at the LGM, their contiguity differed, influencing the spatial distribution of relictual habitat and glacial refugia. For example, Northern and parts of the Central Europe (EU) were largely covered by a single ice sheet (**Fig. 2**) (Hewitt, 1999; Hewitt, 2000). Thus, many high latitude species underwent southward contraction (Hewitt, 2000; Galbreath & Cook, 2004), which result in higher genetic diversity in southern areas. Similarly, in North America (NA), an extensive region was covered by the Laurentide and Cordilleran Ice Sheets (**Fig. 2**) (Hewitt, 2000; Shafer *et al*., 2010) and these regions have species whose genetic diversity decreases with increasing latitude. However, previous studies found that most tree species showed no significant trend (Gougherty, 2022). Furthermore, in Eastern NA, Lumibao *et al*. (2017) showed across multiple species that mid- to high-latitude populations do not share southern haplotypes, and the genetic diversity in mid- to high-latitude is comparable to that in low-latitude, conclusing little support for the LG. Finally, East Asia (EA), though at similar latitudes, was not covered by a unified ice sheet in the LGM, but was rather characterized by multiple scattered glaciers (**Fig. 2**) (Shi, 2002). Many plant species in EA survived in *situ* in multiple refugia at mid to high latitudes during the LGM (**Fig. 1b**) (Qian & Ricklefs, 2001; Rull, 2009), which helped to maintain genetic diversity (Hu *et al*., 2008; Bai *et al*., 2010; Hao *et al*., 2018). A case in point is the Tertiary relict species *Liriodendron chinense*, which was affected by Quaternary climatic fluctuations and split into several refugia, but by the end of the LGM the differences in genetic diversity among these refugia were not significant (Yang *et al*., 2016). Here, we aim to examine the universality of LG and CMH in genetic diversity patterns across the three Northern Hemisphere regions. We hypothesize that differences in ice sheet extent and contiguity at the LGM will influence the strength of the LG and CMH of genetic variation.

**Fig. 2.**
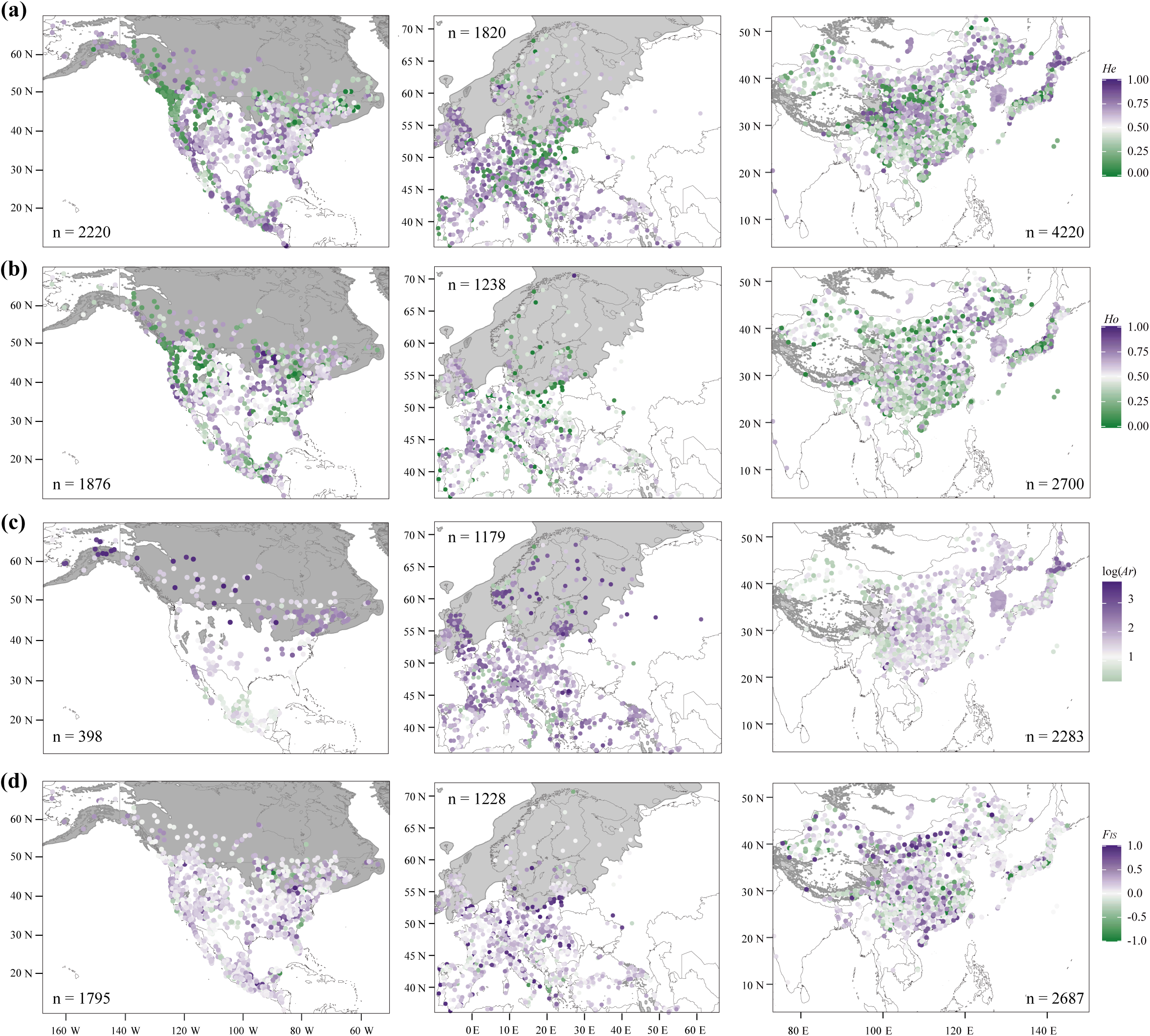
The distribution of genetic variation and ice sheet coverage (gray) in Europe, East Asia, and North America. (a) expected heterozygosity (*H_e_*), (b) observed heterozygosity (*H_o_*), (c) logarithm allelic richness [log (*A_r_*)] and (d) inbreeding coefficient (*F_IS_*). Letter n represents the number of samples. Source data are provided in Table S1.

Complementing these spatial processes, we also expect patterns of genetic variation to be influenced by species-specific life history traits (Hamrick & Godt, 1996; De Kort *et al*., 2021). For example, woody plants may be better able to cope with the impacts of genetic drift in range margin populations due to traits such as high outcrossing rates, long-distance dispersal of pollen and seeds, large effective population size, long lifespans and overlapping generations (Gaut *et al*., 2015; Chung *et al*., 2020). Conversely, herbaceous plants have shorter life cycles and faster reproductive rates, enabling them to more quickly adapt to environmental changes (Hamrick & Godt, 1996). However smaller population sizes make them more susceptible to genetic drift, and their effective population sizes may fluctuate significantly between years, leading to pronounced bottleneck effects (Gaut *et al*., 2015). Consequently, the loss of genetic diversity in expanding leading and range margin populations is expected to be more severe in herbaceous plants.

In this study, using genetic variation data from 445 published plant species, we address three research questions. First, we evaluate evidence for the LG shaping the spatial patterns of contemporary genetic variation in EU, NA and EA. Second, we evaluate evidence for CMH across these same landscapes. In both cases we evaluate alternative drivers, and consider the extent to which these three regions’ contrasting patterns of historic glaciation may explain observed patterns and the potential to generalize genetic diversity patterns across hemispheres. Third, we investigate how species’ life history traits mediate these patterns by comparing evidence for LG and CMH in woody versus herbaceous species. Finally, we illustrate that a clearer understanding of the mechanisms driving landscape patterns of genetic diversity is integral for wide-scale biodiversity management under ongoing climate change and projected range shifts.

## Materials and Methods

### Literature search and data extraction

We searched for journal articles published before September 6, 2022 in Web of Science using the following keywords: (gene*-diversity OR gene*-variation OR gene*-structure OR population-structure OR population-gene* OR molecular variation) AND (China OR Japan OR *Korea OR Mongolia OR East-Asia) AND (microsate* OR *SSR*) NOT (marine OR ocean OR fresh-water) in EA. We retrieved publications from NA and EU using the same keywords, replacing only the region with (North-America OR United-States OR Canada OR Mexico) and (Europe). We obtained 2,241, 2,386 and 3,564 publications in EU, NA and EA, respectively. We also obtained the dataset of 91 plant species from López-Delgado and Meirmans (2022) and 22 plant species from (Roberts & Hamann, 2015) in NA. For comparison, we also used the keywords (single nucleotide polymorphism OR SNP) combined with regional keywords to search for literature. Since the SSRs are more sensitive to recent demographic changes than SNP and the SNP data is limited (only 998 populations across three regions), we focus on SSR data in this study. The SNP results are presented in the appendix.

Then we filtered the articles according to the following criteria: (1) only populations of plant species were included. (2) introduced, invasive, and cultivated species were excluded. (3) populations reporting at least one of the four genetic diversity parameters [expected heterozygosity under the Hardy–Weinberg equilibrium (*H_e_*), observed heterozygosity (*H_o_*), allelic richness (*A_r_*), or inbreeding coefficient (*F_IS_*)] for included for subsequent analyses. (4) all populations have geographical coordinates for spatial analysis. (5) We set a minimum threshold of three populations per species to ensure statistically reliable comparisons. The filtering process for literatures is detailed in **Fig. S1**, with the raw data and literatures provided in **Tables S1 and S2**. We extracted the altitude for each population using SRTM derived elevation data at 30-second resolution (Farr *et al*., 2007; Fick & Hijmans, 2017).

Some species were reported in multiple studies, however due to differences in SSR markers, the data could not be simply combined for further analysis. Therefore, we prioritized studies that provided more comprehensive genetic diversity parameters, or retained the study with a larger number of populations. Overall, 18, 19 and 41 duplicate species were removed in EU, NA and EA respectively (**Table S3**). We produced semi-variograms for all genetic parameters to visually assess evidence of spatial autocorrelation in gstat package (Pebesma, 2004). The results indicated no or only weak spatial autocorrelation for the three regional genetic diversity parameters (**Fig. S2**), and therefore spatial autocorrelation was not included in the subsequent analysis.

### Calculating the extent of glaciation

We used the st_area function in the sf package (Pebesma & Bivand, 2023) to calculate the ratio of glacier area and geographic area for EU, NA and EA, respectively. The results show that glacier coverage was 31% in EU, 46% NA and 3% in EA, with a single ice cap in EU, two large ice caps in NA and numerous small ice caps in EA.

### Predicting species’ range areas

We collated 19 climatic variables for the current (1960–1990) and LGM (c. 21,000 years ago) period from the WorldClim1.4 database (Hijmans *et al*., 2005), together with the terrain roughness index and SAGA-GIS topographic wetness index from the ENVIREM dataset (Title & Bemmels, 2018). The LGM period was estimated by the MIROC-ESM climate model (Watanabe *et al*., 2011). The spatial resolution of climate variables was 2.5 arc-minutes, ∼5km. We evaluated collinearity using a Spearman’s rank correlation threshold of > 0.7, implemented in the virtualspecies R package (Leroy *et al*., 2016), selecting one variable from group of collinear variables. This resulted in five climatic variables as well as one terrain variable in each region: mean diurnal temperature range (Bio2), temperature annual range (Bio7), mean temperature of wettest quarter (Bio8), annual precipitation (Bio12), precipitation seasonality (Bio15), terrain roughness index in the EU and EA and SAGA-GIS topographic wetness index in the NA.

We augmented published population occurrence records with additional records downloaded from Global Biodiversity Information Facility (GBIF, www.gbif.org). The dataset can be accessed via DOI provided in **Table S4**. Each species’ occurrences were cleaned using the CoordinateCleaner package (Zizka *et al*., 2019). A minimum of ten coordinates were required to calibrate a species distribution model (SDM) (Van Proosdij *et al*., 2016). We used Generalized Linear Models (GLM) (McCullagh & Nelder, 1989), Maximum Entropy (MaxEnt) (Phillips *et al*., 2006) and Domain (Carpenter *et al*., 1993) to build an ensemble SDM using the sdm R package (Naimi & Araújo, 2016). Ensembles were generated using weighted averaging based on the AUC statistic. We used 70% of the data for training and the 30% remaining data to evaluate the model and repeated this process ten times. We evaluated the performance of our models using the area under the receiver operating characteristic curve (AUC). We then calculated the SDM threshold based on the maximum test sensitivity and specificity (MaxSSS) (Liu *et al*., 2013), and converted the prediction outputs at the current and LGM into a binary presence-absence map.

### Generating species’ range and niche position

We calculated climatic suitability for each population, under current and LGM climate, as well as distance to the refugia, distance to the range margin and distance to the native climatic niche margin. We converted the LGM binary presence-absence map to spatial presence points, where the presence points represent species refugia. We then calculated the distance from each population to the refugia (i.e., the closest distance of each population to presence points) (**Fig. 1a, b**). We converted the current binary presence-absence map to spatial polygons and then calculated distance to the range margin (i.e., the closest distance of each population to the margin of the polygon) (**Fig. 1d**). For a small number of populations that were located outside of the polygon (resulting from our MaxSSS threshold), we assigned distance as zero, which means that those populations were at the range margin.

We performed principal component analysis (PCA) on all 19 climatic variables from the worldclim1.4 (Hijmans *et al*., 2005) within EU (Longitude: -10 - 66 degree; Latitude: 36 - 72 degree), NA (Longitude: -167 - -50 degree; Latitude: 10 - 80 degree) and EA (Longitude: 73 - 150 degree; Latitude: 4 - 53 degree), using the ENMGadgets R package (Barve & Barve, 2013). Together, the first two axes accounted for over 70% of the climatic variation in each region, so we used PC1 and PC2 to construct the native climatic niche space (CNS) for the species. For each species, we used the filtered occurrence records (see above for details) to extract values from the first two axes of PCA and then used a kernel density estimator in ks R package (Duong, 2007) to extract the contour line corresponding to 99% of the estimated density. We converted the contour lines to polygons, which defined the CNS (**Fig. 1c**). We used the NMI function from Broennimann *et al*. (2021) to calculate the shortest distance to the native climatic niche margin for each population (**Fig. 1c**). NMI stands for Niche Margin Index, which is an indicator used to assess the degree of match between a population’s climatic conditions and its species’ CNS. The main calculation process of the NMI function is (1) assigned a positive sign to NMI if the population sites were located inside the CNS, and a negative sign if outside the CNS. (2) calculated the minimum orthogonal distance of population sites to the CNS margin using the gDistance function of the R package rgeos (Bivand *et al*., 2016). (3) scaled each distance between -1 and 1 (i.e., a value of 1 indicates the population further away from CNS margins). For a small number of populations that were located outside of the modelled CNS, we assigned distance as zero, which means that those populations were at the native climatic niche margin.

### Bayesian path analysis of drivers of genetic variation

To test how contemporary and historical climate, latitude and altitude affect the spatial pattern of population genetic variation, we hypothesized that climate, niche, latitude and altitude have a direct effect on genetic variation, as well as an indirect effect through population geographic location on genetic variation. We represented these relationships using a path diagram (**Fig. S3**). We employed Bayesian phylogenetic structural equation modelling to assess the strength of association. The species names were calibrated using world flora online plant list (wfoplantlist.org, WFO). We used the V.PhyloMaker2 package to obtain species’ phylogenetic placement, which is based on the largest phylogeny of vascular plants (GBOTB.extended.tre) (Jin, 2024). Then, we applied the vcv.phylo function in the ape package to convert the phylogenetic tree into a variance-covariance matrix (Paradis & Schliep, 2019), which was incorporated into the Bayesian model as a random effect. Prior to analyses, we performed a log transformation on the data for genetic parameters (*H_e_*, *H_o_*, *A_r_* and *F_IS_*) to reduce heteroskedasticity (Moles *et al*., 2009), and scaled D_niche_, D_range_, D_ref_, S_cur_, S_LGM_, latitude and altitude to a mean of 0 and a standard deviation of 1, to facilitate the comparison of coefficient estimates (Schielzeth, 2010).

### Comparison of woody and herbaceous plants

Species were classified into woody or herbaceous forms based on Luo *et al*. (2023), Flora of China (www.iplant.cn), Flora of North America (floranorthamerica.org) and European and Mediterranean vascular plant database (europlusmed.org). We ran Bayesian models separately for woody and herbaceous species, with four chains comprising 4000 iterations and 1500 for burn-in. We used weakly informative normal priors on beta parameters with a mean of 0 and standard deviation of 1 and default priors for other model parameters (Schmidt *et al*., 2022). We performed all analyses using the brms package (Bürkner, 2017). The Rhat values for all the models were below 1.01, and the effective sample sizes (ESS) were above 1000, confirming that all chains converged (Gelman *et al*., 2015). The posterior distribution and trace plots of the models are shown in **Fig. S4**. All analyses were performed in R-4.1.1 (R Core Team, 2021).

## Results

We compiled genetic variation data from 402 articles, representing 445 species and 8530 populations (1909 for EU, 2290 for NA and 4331 for EA) (**Fig. 2**, **Table S1 and S5**). Woody plants accounted for 52%, 69% and 74% of species in EU, NA and EA, respectively (mean = 65%). SDMs were developed for 445 species, retaining 10 to 8069 occurrences (mean = 594) per species (**Table S4**). SDMs had a high predictive power with Domain, GLM and Maxent having a minimum AUC value of 0.7 respectively (**Fig. S5**, **Table S4**), therefore we retained all species for subsequent analysis.

### Evidence for the latitudinal gradient hypothesis

We observed significant differences in the LG of genetic diversity across EU, NA, and EA. Specifically, genetic diversity (*H_e_ H_o_* and *A_r_*) was negatively associated with latitude in EU and positively associated in EA, with no significant relationship observed in NA (**Fig. 3**). The *F_IS_* showed no significant relationship with latitude in any of the studied regions (**Fig. 3d, h, l**). Populations in EU (*H_o_* and *A_r_*) and NA (*H_e_*) exhibit significantly higher genetic diversity when further away from refugia, whereas those in EA (*H_o_* and *A_r_*) show lower genetic diversity (**Fig. 3**). Populations located further away from refugia exhibited lower *F_IS_* in EU (**Fig. 3d**).

**Fig. 3.**
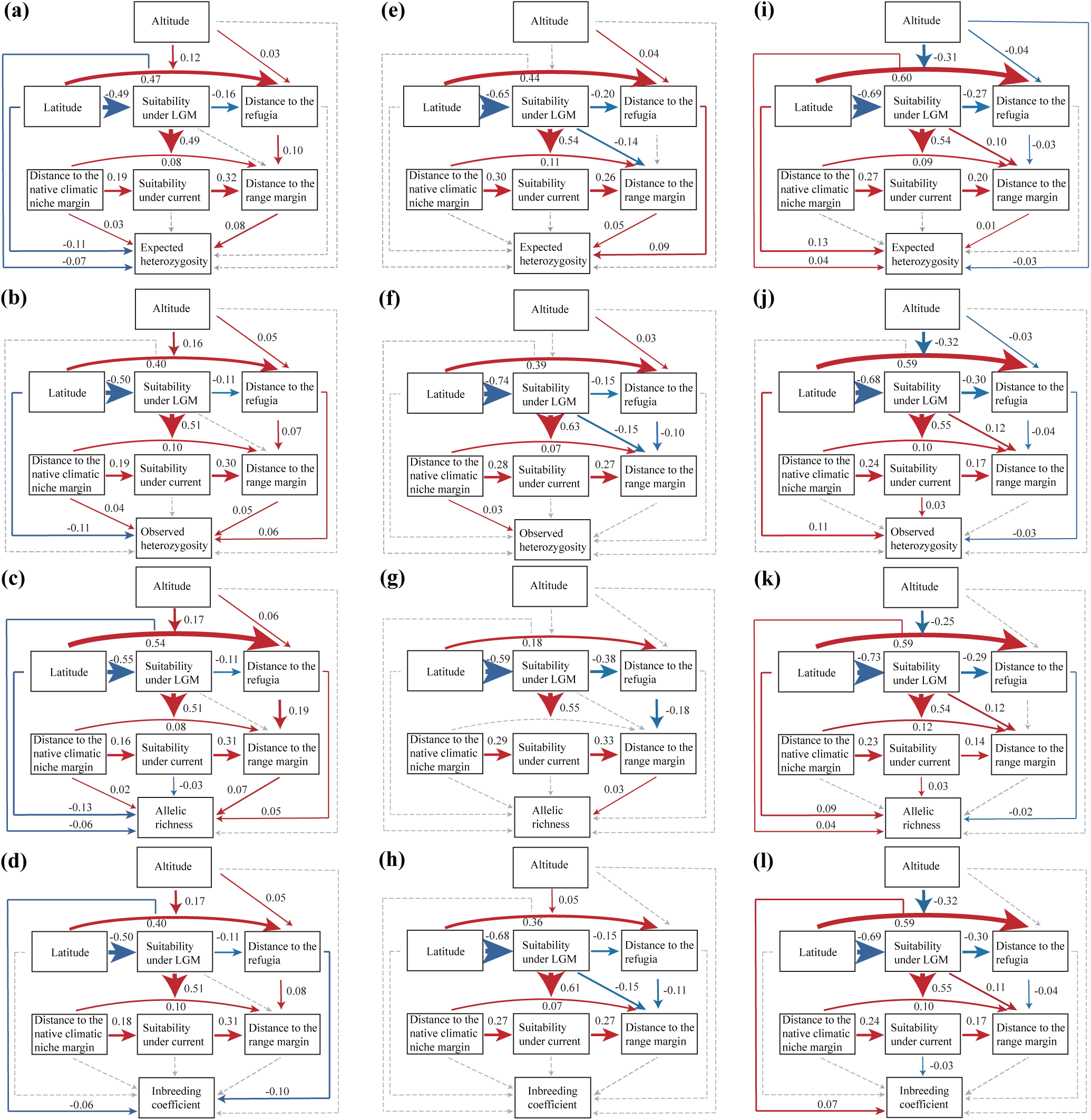
The path model illustrating the direct impacts of biogeographic position, habitat suitability, altitude and latitude on expected heterozygosity (a, e, i), observed heterozygosity (b, f, j), allelic richness (c, g, k) and inbreeding coefficient (d, h, i) across Europe (a, b, c, d), North America (e, f, g, h) and East Asia, as well as the indirect impacts of species’ climatic niche, altitude and latitude on those genetic variation. Blue and red arrows represent negative and positive effects, respectively, with the corresponding mean value on the arrow. Dashed and solid lines represent 95% credible intervals overlapping with zero or not, respectively.

### Evidence for the central-marginal hypothesis

We observed patterns of genetic diversity that conform to the CMH in all regions. In EU (*H_e_*, *H_o_*, and *A_r_*), NA (*H_e_* and *A_r_*) and EA (*H_e_*), populations that are further away from the range and/or niche margins exhibited higher genetic diversity (**Fig. 3**). Other genetic diversity indicators in these regions were non-significant, but none showed a negative relationship. The *F_IS_* showed no significant relationship with range and niche positions in EU, NA and EA (**Fig. 3d, h, l**).

### Comparison of woody and herbaceous plants

The LG for woody and herbaceous plants was highly variable. Woody plants are negatively correlated with latitude in NA, positively correlated in EA, and uncorrelated in EU (**Fig. 4b [Path 2]**). The LG for herbaceous plants were negatively correlated with latitude in EU and positively correlated in NA and EA (**Fig. 4b [Path 2]**). The genetic diversity patterns of woody plants align with the CMH in all regions (**Fig. 4b [Path 3 and 4]**). Genetic diversity of herbaceous plants was also positively correlated with distance to range and niche margin in EU, but non-significant in EA **and NA (**Fig. 4b [Path 3 and 4]).

**Fig. 4.**
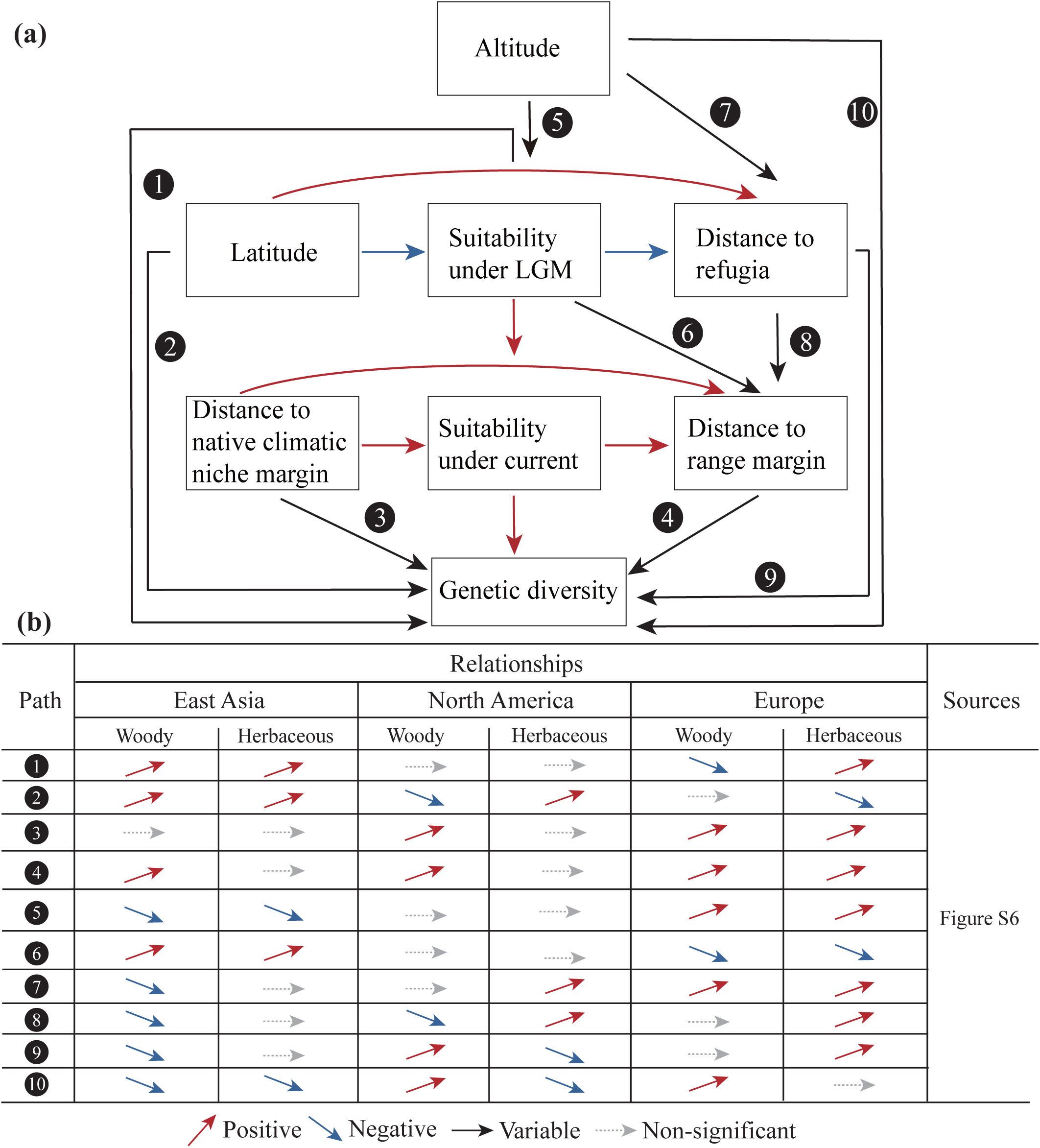
The pathway concept diagram demonstrating the direct and indirect relationships between ecological variables and genetic diversity for woody and herbaceous plants acoss Europe, East Asia and North America. The blue, red, and black arrows represent the positive, negative, and variable correlations of the path diagram (a) Detailed descriptions of the change pathways within the datasets of woody and herbaceous plants (b) These relationships summarize the results shown in Fig. S6.

### Comparison of SSR and SNP datasets

SNP data literature screening and retention are shown in **Fig. S7**. SNPs are consistent with SSRs in showing non-significant correlations between genetic diversity and latitude in EU and NA and inconsistent in EA where SNPs show a negative correlation whereas SSRs show a positive correlation (**Fig. S8**). SNPs do not support the CMH whereas SSPs support the CMH (**Fig. S8**).

## Discussion

Based on the 445 species, we find that LG pattern of genetic diversity occur inconsistently across EU, EA and NA in the Northern Hemisphere. By contrast, we find broad and consistent evidence for the CMH. The CMH is stronger in woody than in herbaceous plants. Below we discuss the patterns of genetic diversity in the context of latitude and population position in detail. In addition, we discussed the impact of life history in shaping the patterns of genetic diversity.

### Patterns of genetic diversity along latitudinal gradients shaped by glaciers

The differences in glaciation between EU, NA, and EA have shaped the LG of genetic diversity in each region (Fonseca *et al*., 2023). Genetic diversity was negatively correlated with latitude in EU, uncorrelated with latitude in NA, and positively correlated with latitude in EA (**Fig. 3**). We found that although the high and partly mid-latitude regions of EU and NA are covered by ice sheets, species are contracting along different routes. Northern EU has no major refugia, and species mainly retreated to the peninsulas of Iberia, Italy and the Balkans (Hewitt, 2011). NA western had two macro-refugia: the Beringia in the north and the Pacific Northwest in the south (Hultén, 1937; Pielou, 1991; Shafer *et al*., 2010). In addition, the unglaciated eastern NA has no clear phylogeographical spatial pattern (Soltis *et al*., 2006; Lumibao *et al*., 2017). Therefore, species may have retreated north, south or in both directions during the glacial period, which contributes to the sustainment of genetic diversity in the southern and northern regions. The major refugia of EA plants are widely distributed in different latitudes, such as the Changbai Mountains in Northeast, the Qinling Mountains in central, the Hengduan Mountains in Southwest and the Tianshan Mountains in Northwest (Qiu *et al*., 2011; Liu *et al*., 2012; Tang *et al*., 2018). Thus, during glaciation, EA species contracted in *situ* to multiple refugia (Hu *et al*., 2008; Bai *et al*., 2010; Hao *et al*., 2018; Xu *et al*., 2021). The evidence from EA suggests that local expansion of species occurred after the glacial period, i.e., populations moved closer to refugia while also closer to range centres (**Fig. 3i, j, l**).

Thus, species at higher latitudes are able to occupy more range more rapidly than species at lower latitudes and therefore may have larger mean effective population sizes and hence higher genetic diversity (Fine, 2015; Bernos & Fraser, 2016; Lawrence & Fraser, 2020). Thus, local contraction and expansion along the LG have shaped the pattern of genetic diversity in EA. In addition, our results also show that populations further from the Himalayan glaciers exhibit higher genetic diversity (*H_e_* and *H_o_*) (**Table S7**), suggesting the presence of an east-west genetic diversity pattern in EA. The east-west mountain ranges in EA not only promote the formation of micro-refugia but also serve as corridors for species dispersal (Tian *et al*., 2018; Zhang *et al*., 2018). The pattern is also evidenced by the negative relationship between altitude and genetic diversity in EA (**Fig. 3i**).

### Patterns of genetic diversity are consistent with the central-marginal hypothesis across the Northern Hemisphere

We find evidence for the CMH across all regions, with the strongest relationship in EU (**Fig. 3a, b, c**). In EU, species expanded northward from multiple refugia in the peninsulas of Iberia, Italy and the Balkans (Gómez & Lunt, 2007), and different lineages admixed at mid-latitudes (Petit *et al*., 2003), evolving into centres of genetic diversity, and the refugia became the margins (Pironon *et al*., 2017). The evidence from EU is consistent with this pattern, i.e., populations further away from refugia have higher genetic diversity (*H_o_* and *A_r_*), while populations located in refugia are also located at range margins (**Fig. 3b, c**). Further expansion of the admixed populations, combined with a decrease in genetic diversity of leading-edge populations due to founding events and bottleneck effects generated contemporary observed patterns. We find similar, but slightly weaker patterns in NA and EA. The lack of established migration routes and lineages admixed in the unglaciated regions of NA and EA may be a reason for the weak pattern (Soltis *et al*., 2006; Qiu *et al*., 2011; Liu *et al*., 2012; Lumibao *et al*., 2017). Furthermore, the extensive mountainous ranges in EA and NA may harbor suitable local habitats even at the range margins, thereby weakening the CMH (Pironon *et al*., 2017).

### Life history shapes patterns of genetic diversity

While both woody and herbaceous plants were positively associated with latitude in EA, their trends were opposing in NA and negative for herbaceous plants in EU. In NA, woody plants migrated southward during the glacial period and expanded northward after the postglacial period (Roberts & Hamann, 2015), leading to low genetic diversity at high latitudes. In contrast, most herbaceous plants in NA migrated to Beringia, where genetic diversity was maintained at high latitudes during the glacial (Shafer *et al*., 2010; Hoffecker *et al*., 2014). The opposing LG patterns of woody and herbaceous plants may contribute to the overall weak latitudinal gradient in NA. Genetic diversity of woody plants fits the CMH across EU, EA and NA. For herbaceous plants, we found significant support for the CMH in EU, but no significant pattern in other regions (**Fig. 4b [Path 3 and 4]**). Considering possible explanations for this observation, we highlight that the rate of native climatic niche evolution is reported to be lower in woody plants than in herbaceous plants (Smith *et al*., 2010), and therefore, niche conservatism may restrict the ability of woody species to adapt to new environments perhaps attenuating range edge genetic diversity (Hawkins *et al*., 2011). Similarly long-distance dispersal of pollen and seed spanning several kilometers have been increasingly evidenced in woody plants, potentially generating leading edge bottlenecks as ranges shift (Godoy & Jordano, 2001; Gaiotto *et al*., 2003; Bacles *et al*., 2006; Petit & Hampe, 2006). By comparison, weaker CMH patterns in herbaceous plants may be attributed to several possible explanations.

Foremost we suggest that faster generation times may facilitate a more rapid recovery of diversity (Wei *et al*., 2023), and for a given refugial area, may support a larger effective population size, potentially maintaining higher standing genetic diversity.

### Limitations

To enable comparisons across regions, despite differences in study design and species we implemented strict sampling criteria, integrated a phylogenetic covariance matrix and sought to capture mediator variables as well as direct drivers in our model.

However certain limitations persist on our experiemntal design. For example, some species are sampled without covering their entire range which may hinder assessment of the overall spatial genetic structure. Secondly, this study used SDM to calculate species ranges and refugia. The limitations of SDMs include assumptions that species ranges are directly influenced by environmental factors without fully accounting for biological characteristics or interspecies interactions, which may also bias analysis of species’ genetic diversity patterns. However, given that our literature survey collated basic genetic diversity indicators from studies with a very broad range of aims, the sample size is large, and that the approach was consistent between regions, we feel satisfied that the risk of a bias to impact our overall conclusions is low. We highlight also that analyses of comparative patterns in SNPs, which may be informative about deeper time due to lower mutations rates (Borrell *et al*., 2018), are hindered by the lower data availability, though this will likely improve. Therefore, in future studies, more comprehensive sampling, larger sample sizes and access to a broader range of genetic variations through genome sequencing will give us greater ability to detect weaker signals and analyse more specific traits across narrower groups of species.

### Conclusion

This study seeks to examine the ubiquity of broad scale patterns in genetic diversity across Northern Hemisphere plants. Significant differences in the LG were observed among the three regions, but evidence for the CMH was largely consistent. We suggest that variability in these patterns across Northern Hemisphere landscapes, may be associated with differences in species contraction and expansion induced by glaciation and historical climate change. Current global climate warming trends are likely to further reshape contemporary and future patterns of genetic diversity, and interact with anthropogenic range fragmentation and reduced effective population size. These competing drivers may interact with LG and CMH patterns in new, potentially less predictable ways. These data illustrate how macroecological analyses can address fundamental questions in evolutionary biology and inform contemporary monitoring and management of landscape genetic diversity.

## Acknowledgements

This work was funded by the Youth Innovation Team Project for Talent Introduction and Cultivation in University of Shandong Province to Nian Wang and the Outstanding Young Scholars selected by the State Forestry and Grassland Administration, China and the Taishan Scholars Program to Dafeng Chen.

## Author contributions

LL, JSB and NW designed the research. LL collected the data and performed the data analysis. LL, JSB and NW wrote the manuscript.

## Data availability

All study data and the scripts for data processing and analysis are available in the article and Supporting Information.

