## Supplementary material for "Differing drivers of range-wide genetic diversity across previously glaciated Northern hemisphere landscapes": Supporting Information20250428.docx

**Supporting Information:**

Methods based on SNP Data: We used the keywords (single nucleotide polymorphism OR SNP), (China OR Japan OR *Korea OR Mongolia OR East-Asia), (North-America OR United-States OR Canada OR Mexico), and (Europe) to separately search the relevant literature for EA, NA, and EU in Web of Science. We obtained 3,564, 2,386 and 2241 publications in EA, NA and EU, respectively. The criteria for filtering and retaining literature were consistent with those for SSR, with the addition of nucleotide diversity as a parameter for genetic diversity. The filtering process for literature is detailed in Fig. S7, with the raw data and literatures provided in Tables S1 and S2. In both EU and EA, there was one species sourced from different studies. We removed the study with fewer populations for each case (Table S3). Finally, we compiled a genetic variation dataset for 10 articles representing 10 species and 197 populations in EU, 7 articles representing 9 species and 131 populations in NA and 36 articles representing 42 species and 670 populations in EA (Tables S1 and S6). SDMs were developed for 61 species in EU, NA and EA, retaining 15 to 4609 occurrences retained for each species, with an average of 678 occurrences (Table S4). SDMs had a high predictive power with Domain, GLM and Maxent having a minimum mean AUC value of 0.80 (Table S4), therefore we retained all species for subsequent analysis.

**Figure legends**

**Fig. S1** PRISMA diagram based on SSR. PRISMA (Preferred Reporting Items for Systematic Reviews and Meta-Analyses) flow chart showing the procedure of selecting publications for microsatellite data.

**Fig. S2** Semivariance plot of genetic variation for Europe (a), North America (b) and East Asia (c). The x-axis represents geographical distance, and the y-axis shows semivariance values calculated from the Box-Cox transformed genetic variation.

**Fig. S3** The conceptual framework illustrates the direct impacts of biogeographic position, habitat suitability, altitude and latitude on genetic diversity for plants species, as well as the indirect effects of species' climatic niche, altitude and latitude on those genetic diversity. Blue, red and black arrows represent negative, positive and variable effects, respectively.

**Fig. S4** Posterior distributions and trace plots of the Bayesian model for genetic variation and variables in Europe (blue), North America (orange) and East Asia (green). The posterior distributions of the parameters illustrate the estimated values of key parameters and their uncertainties, while the trace plots demonstrate the convergence and stability of MCMC sampling. The model was constructed using brms. The trace plots show no apparent trends, indicating good convergence of the model.

**Fig.S5** Frequency distribution histogram of area under curve (AUC) for species distribution modeling. Source data are provided as Table S4.

**Fig. S6** The path model illustrates the direct impacts of biogeographic position, habitat suitability, altitude and latitude on expected heterozygosity, observed heterozygosity, allelic richness and inbreeding coefficient for Europe (a, b, c, d), North America (i, j, k, l), and East Asia woody (q, r, s, k) plants and Europe (e, f, g, h), North America (m, n, o, p) and East Asia herbaceous (u, v, w, x) plants as well as the indirect effects of species' climatic niche, altitude and latitude on those genetic variation. Blue and red arrows represent negative and positive effects, respectively, with the corresponding mean value on the arrow. Dashed and solid lines represent 95% credible intervals overlapping with zero or not, respectively.

**Fig. S7** PRISMA diagram based on SNP. PRISMA (Preferred Reporting Items for Systematic Reviews and Meta-Analyses) flow chart showing the procedure of selecting publications.

**Fig. S8** The path model illustrates the direct impacts of biogeographic position, habitat suitability, altitude and latitude on expected heterozygosity, observed heterozygosity, nucleotide diversity and inbreeding coefficient for Europe (a, b, c), North America (d, e, f, g) and East Asia (h, i, j, k) dataset based on SNP, as well as the indirect effects of species' climatic niche, altitude and latitude on those genetic variation. Blue and red arrows represent negative and positive effects, respectively, with the corresponding mean value on the arrow. Dashed and solid lines represent 95% credible intervals overlapping with zero or not, respectively.

**Table legends**

**Table S1** Genetic summary estimates of all populations in East Asia (EA) and North America (NA) and Europe (EU). Columns correspond to molecular marker type [microsatellite markers (SSR) and single nucleotide polymorphism(SNP)]; region coding [East Asia (EA) and North America (NA) and Europe (EU)]; reference number; Species name, genus name, and family name were calibrated by the World Flora Online Plant List; whether or not the species is a woody plant (1 for yes, 0 for no); longitude; latitude; elevation; values for allelic richness (*A_r_*), observed heterozygosity (*H_o_*), expected heterozygosity (*H_e_*), inbreeding coefficient (*F_IS_*) and nucleotide diversity (*π*); suitability values for current; suitability values for LGM; distance to range margins (in km); distance to climatic niche margins (Standardised to between 0 and 1); distance to refugia (in km) and distance from population to Himalayan glaciers (DHG).

**Table S2** Original genetic variation data sources corresponding to the literature.

**Table S3** Removed duplicate species for SSR and SNP.

**Table S4** Information on the use of occurrence data for each species in East Asia (EA) and North America (NA) and Europe (EU) and assessment of SDM accuracy. Columns correspond to molecular marker type [microsatellite markers (SSR) and single nucleotide polymorphism (SNP)]; region coding [East Asia (EA), North America (NA) and Europe (EU)]; species name; the GBIF DOI for downloading species occurrences; species unique occurrences; the specificity and sensitivity maximisation threshold for each species; the AUC values for each model algorithm employed and mean AUC values and references.

**Table S5** Overview of genetic variation [expected heterozygosity (*H_e_*), observed heterozygosity (*H_o_*), allelic richness (*A_r_*) and inbreeding coefficient (*F_IS_*)] dataset in East Asia, North America and Europe for SSR.

**Table S6** Overview of genetic variation [expected heterozygosity (*H_e_*), observed heterozygosity (*H_o_*), inbreeding coefficient (*F_IS_*) and nucleotide polymorphism (*π*)] dataset in East Asia, North America and Europe for SNP.

**Table S7** The relationship between population genetic variation and Himalayan glaciers distance (DHG).

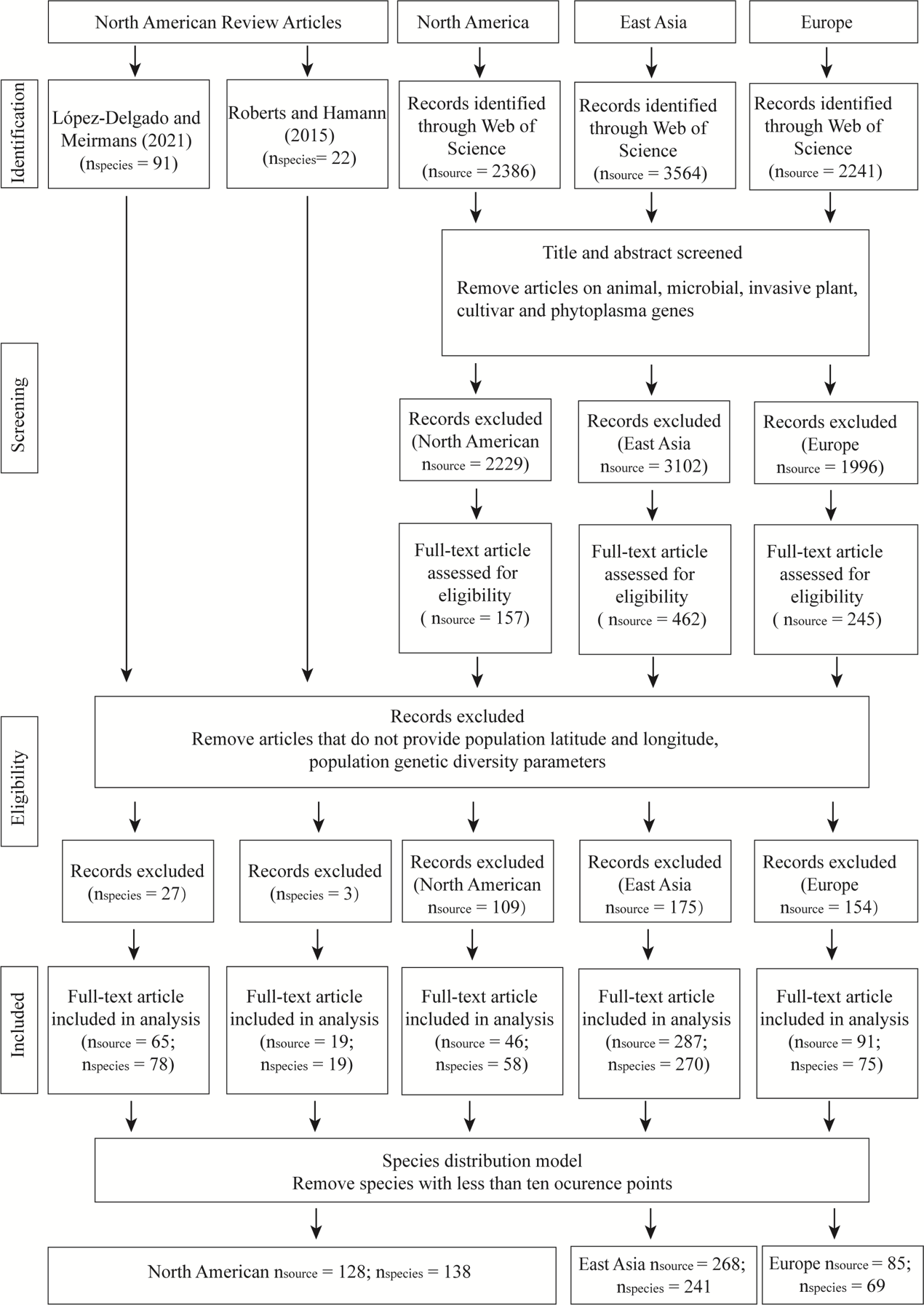
**Fig. S1** PRISMA diagram based on SSR. PRISMA (Preferred Reporting Items for Systematic Reviews and Meta-Analyses) flow chart showing the procedure of selecting publications for microsatellite data.

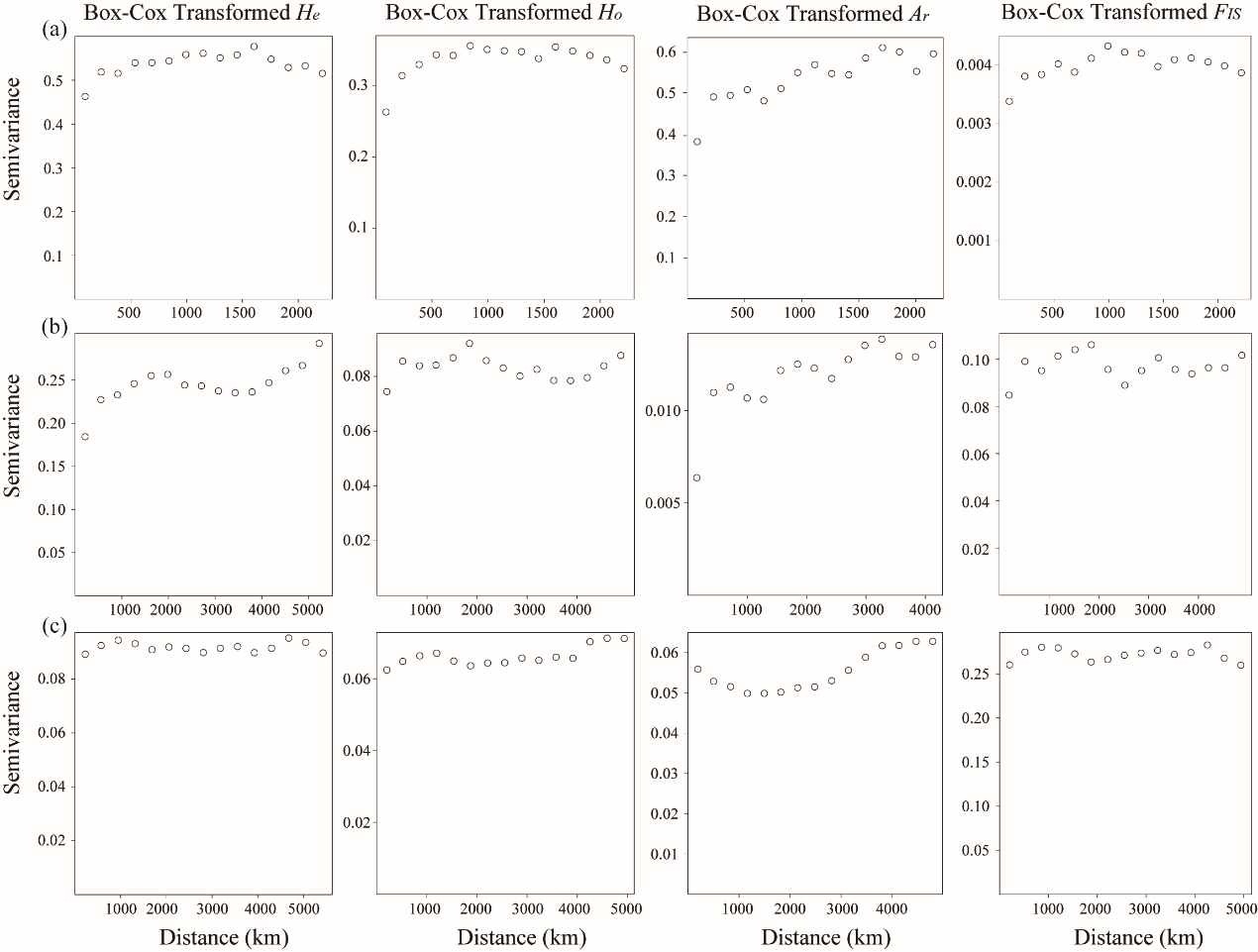
**Fig. S2** Semivariance plot of genetic variation for Europe (a), North America (b) and East Asia (c). The x-axis represents geographical distance, and the y-axis shows semivariance values calculated from the Box-Cox transformed genetic variation.

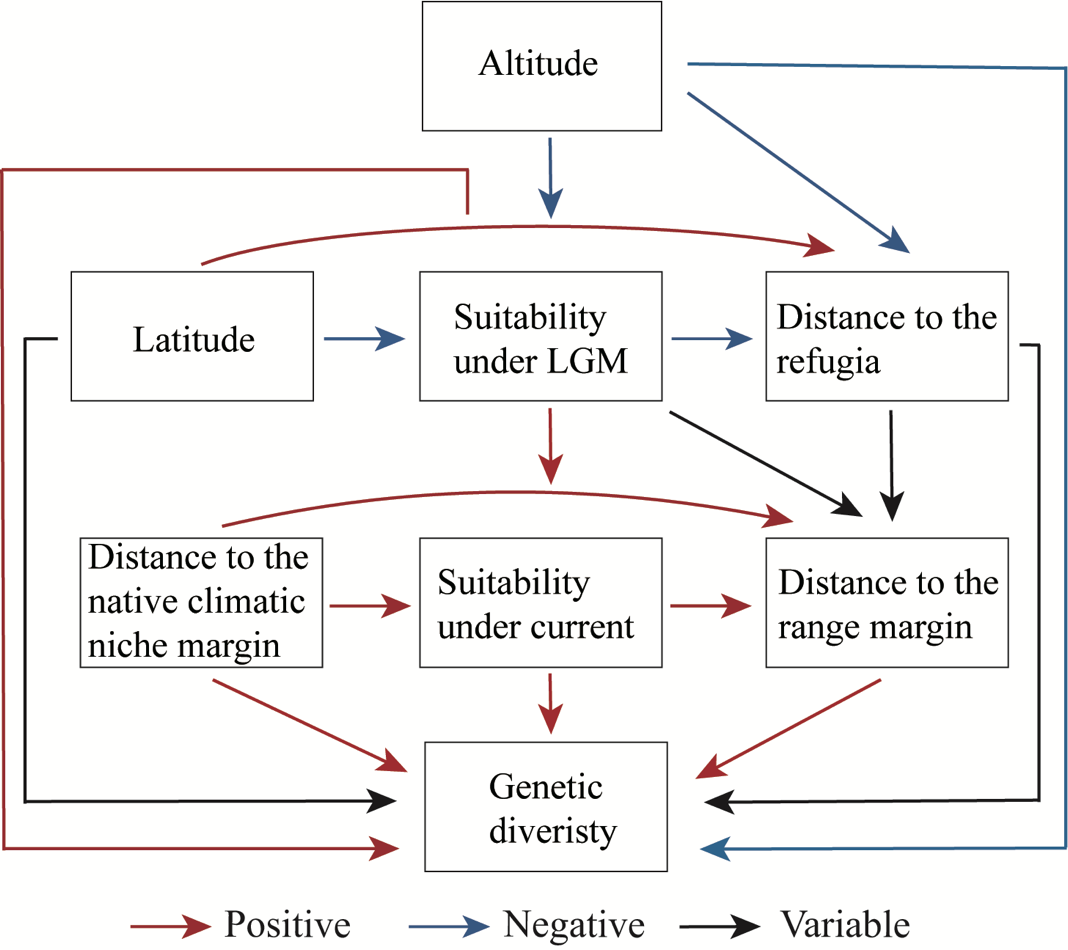
**Fig. S3** The conceptual framework illustrates the direct impacts of biogeographic position, habitat suitability, altitude and latitude on genetic diversity for plants species, as well as the indirect effects of species' climatic niche, altitude and latitude on those genetic diversity. Blue, red and black arrows represent negative, positive and variable effects, respectively.

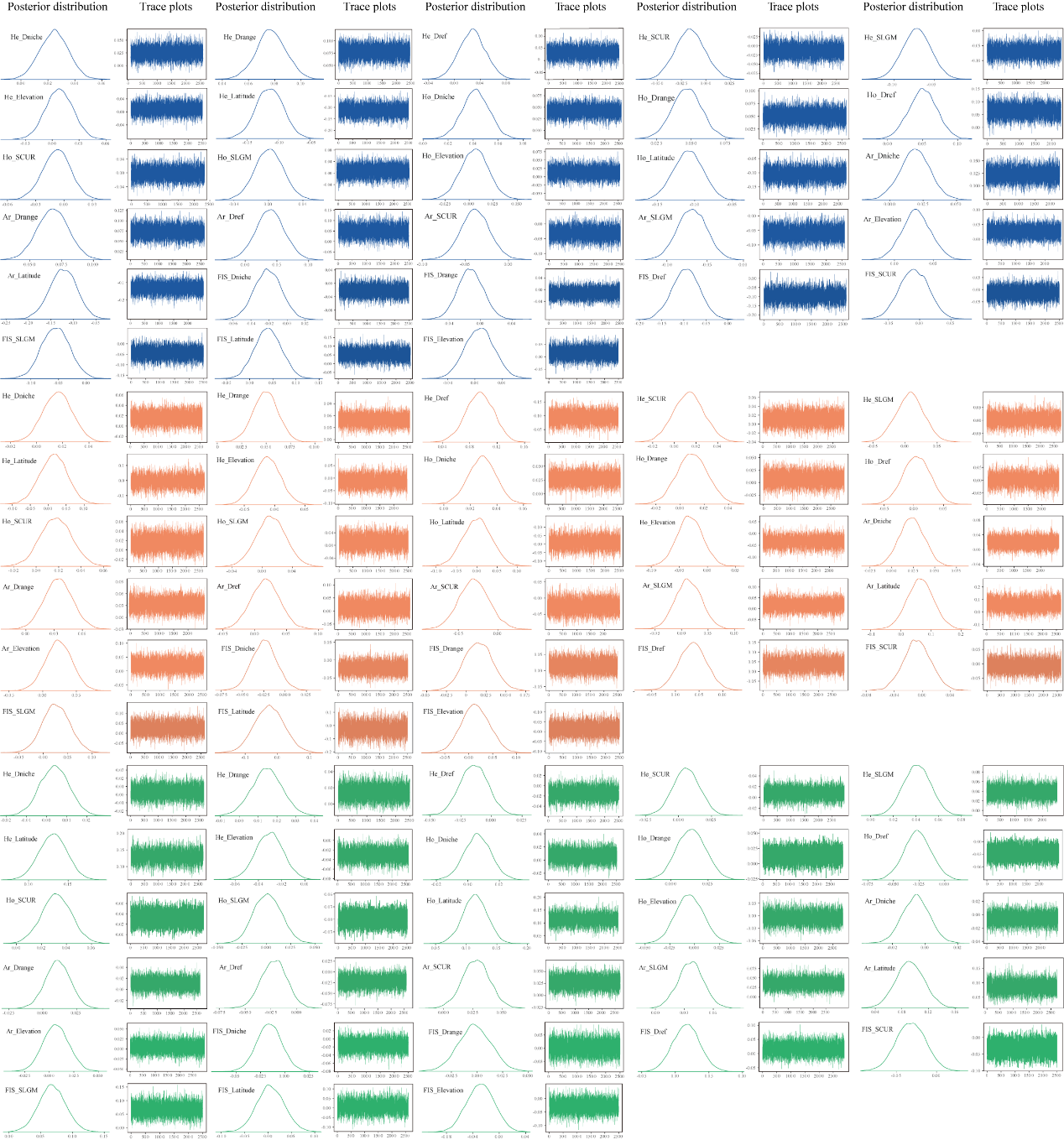
**Fig. S4** Posterior distributions and trace plots of the Bayesian model for genetic variation and variables in Europe (blue), North America (orange) and East Asia (green). The posterior distributions of the parameters illustrate the estimated values of key parameters and their uncertainties, while the trace plots demonstrate the convergence and stability of MCMC sampling. The model was constructed using brms. The trace plots show no apparent trends, indicating good convergence of the model.

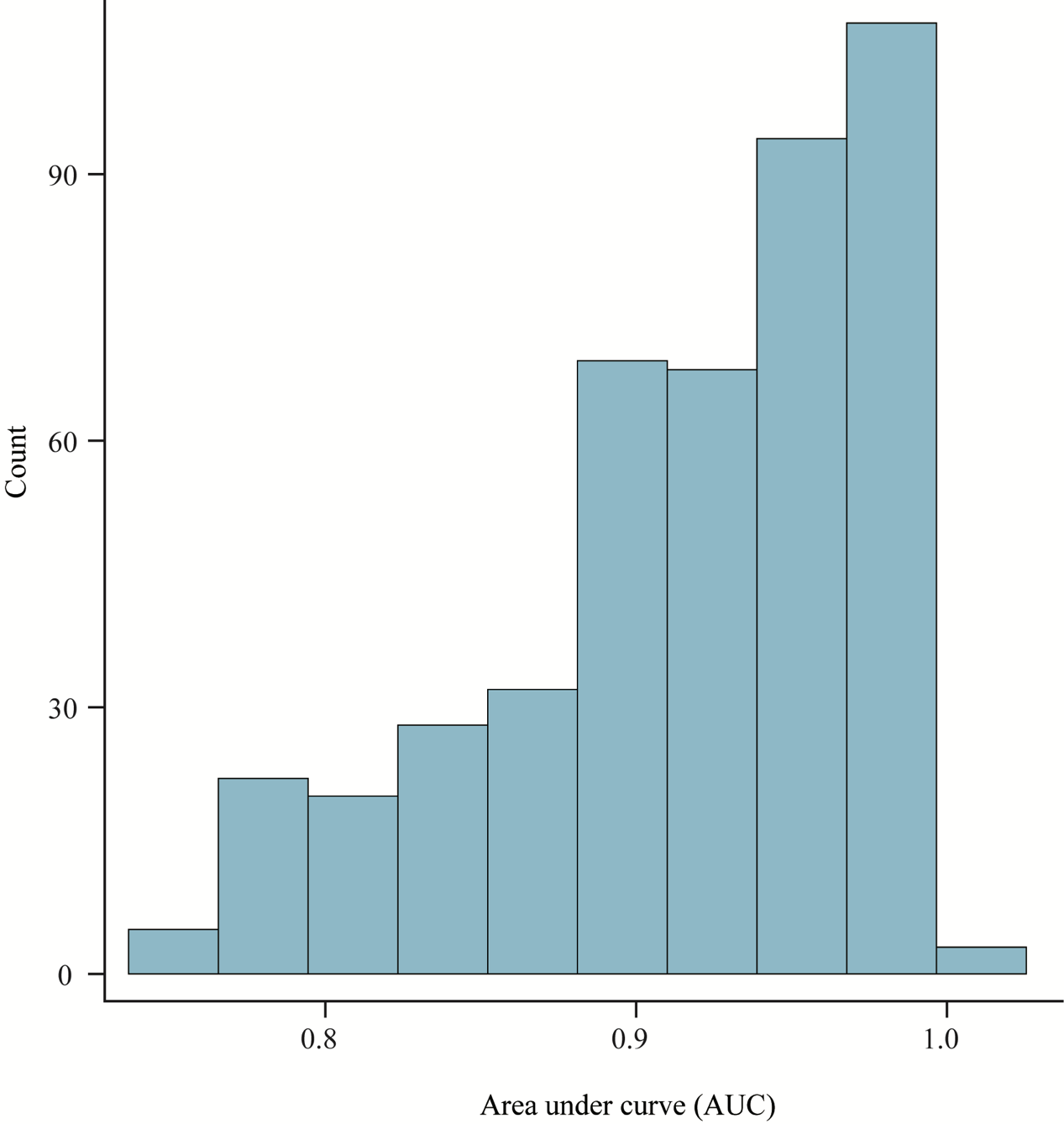
**Fig. S5** Frequency distribution histogram of area under curve (AUC) for species distribution modeling. Source data are provided as Table S4.

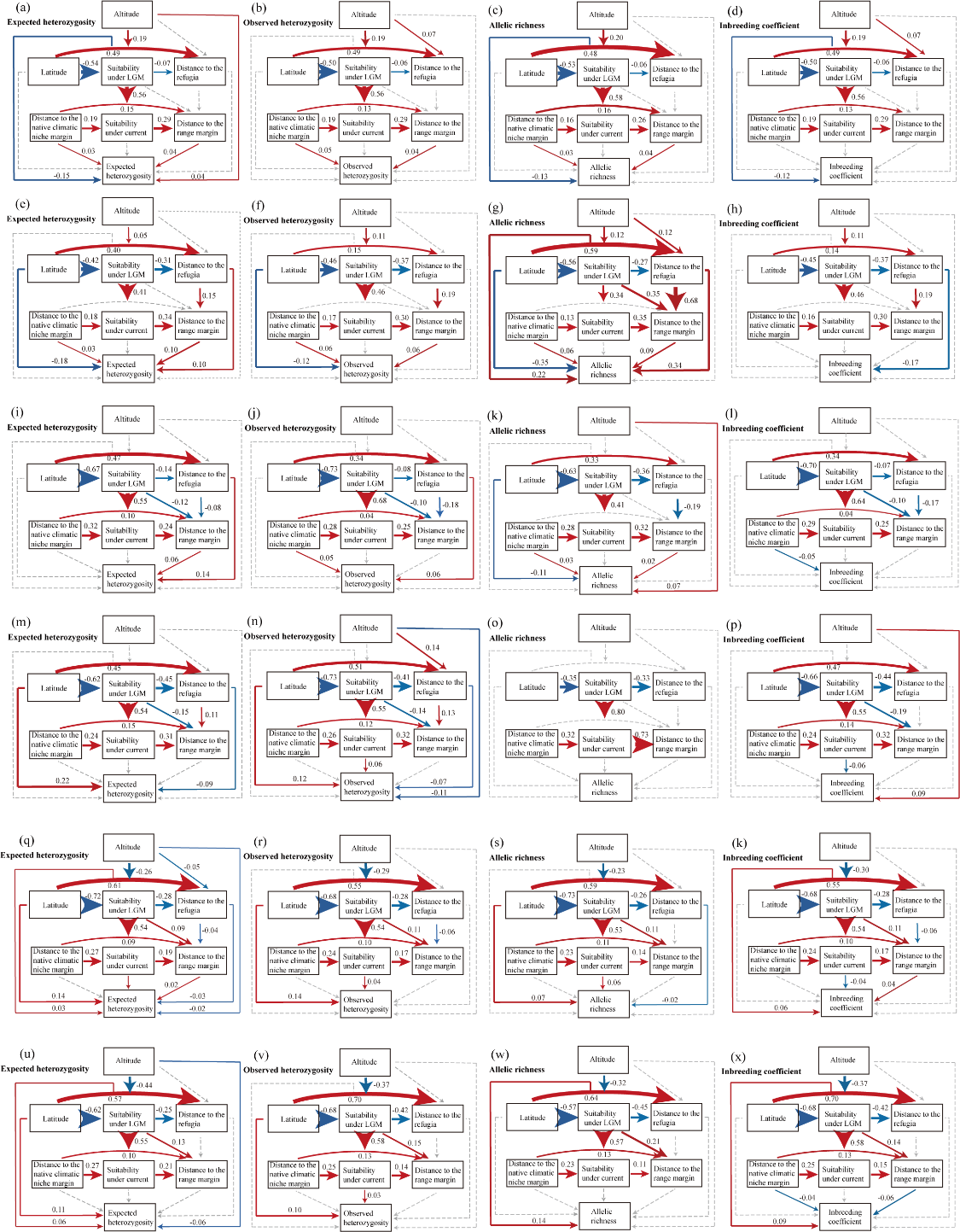
**Fig. S6** The path model illustrates the direct impacts of biogeographic position, habitat suitability, altitude and latitude on expected heterozygosity, observed heterozygosity, allelic richness and inbreeding coefficient for Europe (a, b, c, d), North America (i, j, k, l), and East Asia woody (q, r, s, k) plants and Europe (e, f, g, h), North America (m, n, o, p) and East Asia herbaceous (u, v, w, x) plants. as well as the indirect effects of species' climatic niche, altitude and latitude on those genetic variation. Blue and red arrows represent negative and positive effects, respectively, with the corresponding mean value on the arrow. Dashed and solid lines represent 95% credible intervals overlapping with zero or not, respectively.

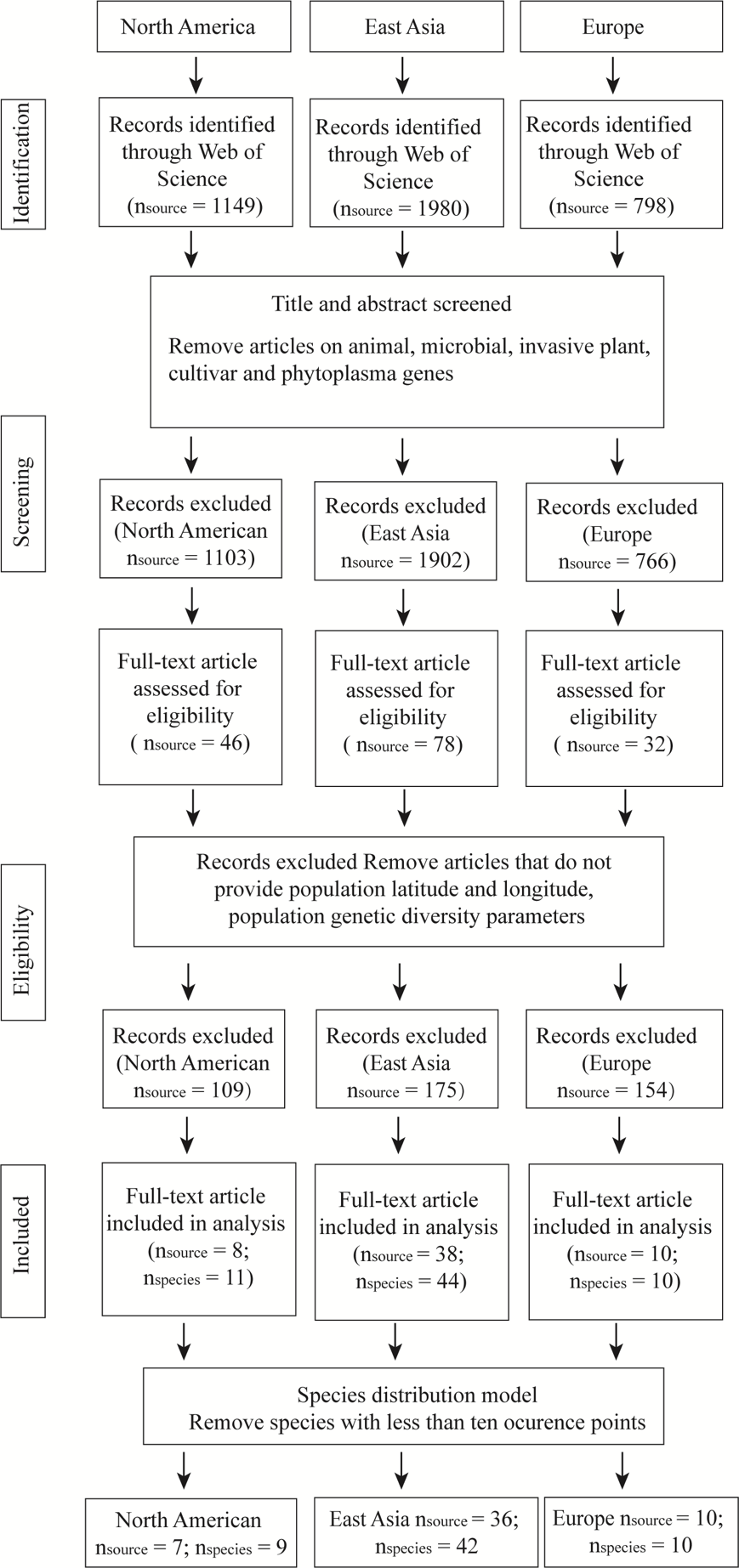
**Fig. S7** PRISMA diagram based on SNP. PRISMA (Preferred Reporting Items for Systematic Reviews and Meta-Analyses) flow chart showing the procedure of selecting publications.

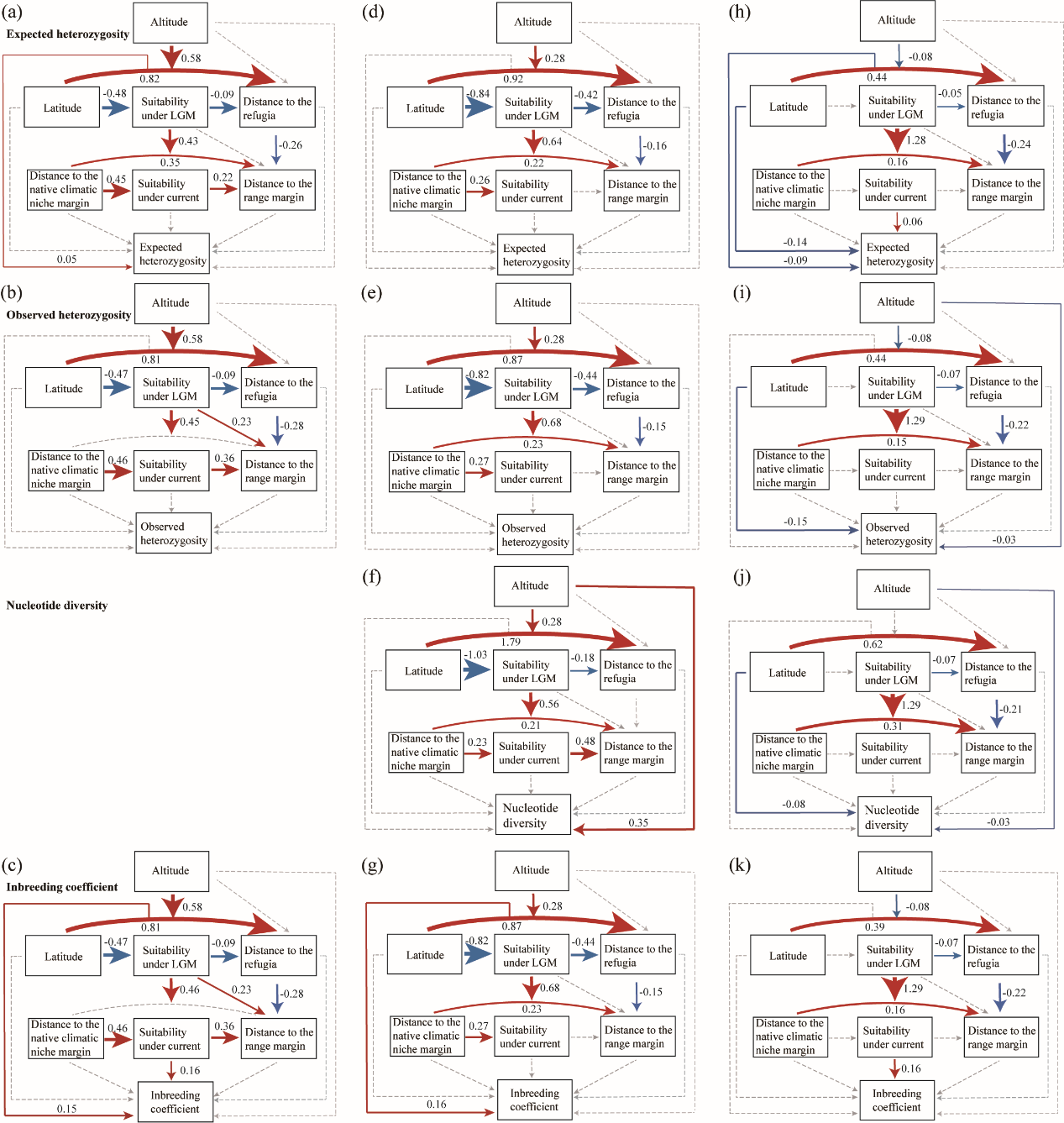
**Fig. S8** The path model illustrates the direct impacts of biogeographic position, habitat suitability, altitude and latitude on expected heterozygosity, observed heterozygosity, nucleotide diversity and inbreeding coefficient for Europe (a, b, c), North America (d, e, f, g) and East Asia (h, i, j, k) dataset based on SNP, as well as the indirect effects of species' climatic niche, altitude and latitude on those genetic variation. Blue and red arrows represent negative and positive effects, respectively, with the corresponding mean value on the arrow. Dashed and solid lines represent 95% credible intervals overlapping with zero or not, respectively.

**Table S5** Overview of genetic variation [expected heterozygosity (*H_e_*), observed heterozygosity (*H_o_*), allelic richness (*A_r_*) and inbreeding coefficient (*F_IS_*)] dataset in East Asia, North America and Europe for SSR.

| **Flora** | **Genetic parameters** | **Number of species** | | **Number of populations** | | **Range of values** | |
| --- | --- | --- | --- | --- | --- | --- | --- |
|  |  | Woody | Herbs | Woody | Herbs | Woody | Herbs |
| East Asia | *H_e_* | 180 | 60 | 3077 | 1058 | 0-0.9 | 0-0.9 |
|  | *H_o_* | 129 | 39 | 2315 | 764 | 0-0.9 | 0-1.0 |
|  | *A_r_* | 89 | 25 | 1728 | 551 | 1.0 - 58.1 | 1.0 - 35 |
|  | *F_IS_* | 128 | 39 | 2303 | 758 | -1.0 - 1.0 | -1.0 - 1.0 |
| North America | *H_e_* | 88 | 45 | 1500 | 663 | 0 - 1.0 | 0 - 0.9 |
|  | *H_o_* | 74 | 45 | 1166 | 672 | 0 - 1.0 | 0 - 1.0 |
|  | *A_r_* | 15 | 4 | 285 | 97 | 1.2 - 35.4 | 1.9 - 8.5 |
|  | *F_IS_* | 71 | 43 | 1131 | 631 | -1.0 - 1.0 | -1.0 - 1.0 |
| Europe | *H_e_* | 32 | 37 | 896 | 866 | 0.1 - 0.9 | 0 - 1.0 |
|  | *H_o_* | 29 | 15 | 832 | 362 | 0.2 - 0.8 | 0 - 1.0 |
|  | *A_r_* | 25 | 14 | 795 | 336 | 1.5 - 10.7 | 1.0 - 8.1 |
|  | *F_IS_* | 28 | 15 | 826 | 359 | -0.8 - 0.8 | -0.5 - 1.0 |

**Table S6** Overview of genetic variation [expected heterozygosity (He), observed heterozygosity (Ho), inbreeding coefficient (FIS) and nucleotide polymorphism (π)] dataset in East Asia, North America and Europe for SNP.

| **Flora** | **Genetic parameters** | **Number of species** | **Number of populations** | **Range of values** |
| --- | --- | --- | --- | --- |
| East Asia | π | 22 | 270 | 0 - 0.6 |
|  | *H_e_* | 41 | 642 | 0 - 0.7 |
|  | *H_o_* | 30 | 517 | 0 - 0.6 |
|  | *F_IS_* | 32 | 547 | -1.0-0.6 |
| North America | π | 5 | 70 | 0.2-0.4 |
|  | *H_e_* | 9 | 126 | 0.03-0.4 |
|  | *H_o_* | 8 | 109 | 0.02-0.3 |
|  | *F_IS_* | 8 | 109 | -0.7-0.8 |
| Europe | π | - | - | - |
|  | *H_e_* | 10 | 180 | 0.02-0.3 |
|  | *H_o_* | 9 | 172 | 0.01-0.3 |
|  | *F_IS_* | 9 | 172 | -0.5-0.5 |

**Table S7** The relationship between population genetic variation and Himalayan glaciers distance (DHG).

| Type | Variable | Coefficient | CIlo95 | CIup95 | CIlo90 | CIup90 | Response_var |
| --- | --- | --- | --- | --- | --- | --- | --- |
| SSR | DHG | 0.07 | 0.04 | 0.09 | 0.04 | 0.09 | *He* |
|  | DHG | 0.06 | 0.03 | 0.09 | 0.04 | 0.09 | *Ho* |
|  | DHG | 0.03 | -1.99E-04 | 0.07 | 0.00 | 0.06 | *Ar* |
|  | DHG | -0.03 | -0.07 | 0.01 | -0.07 | 0.01 | *FIS* |
| SNP | DHG | -0.05 | -0.10 | 0.01 | -0.09 | 0.00 | *He* |
|  | DHG | -0.06 | -0.11 | -0.01 | -0.11 | -0.02 | *Ho* |
|  | DHG | -0.06 | -0.21 | 0.10 | -0.19 | 0.07 | *Fis* |
|  | DHG | -0.04 | -0.08 | -0.01 | -0.08 | -0.01 | *π* |
